# Co-occurrence history increases ecosystem temporal stability and recovery from a flood in experimental plant communities

**DOI:** 10.1101/262337

**Authors:** Sofia J. van Moorsel, Terhi Hahl, Owen L. Petchey, Anne Ebeling, Nico Eisenhauer, Bernhard Schmid, Cameron Wagg

## Abstract

Understanding factors that increase ecosystem stability is critical in the face of environmental change. Experiments simulating species loss from grassland ecosystems have shown that losing biodiversity decreases the ability of ecosystems to buffer negative effects of disturbances. However, as the originally sown experimental communities with reduced biodiversity develop, plant evolutionary processes or the assembly of interacting soil organisms may allow them to develop stability and resilience over time. We explored such effects in a long-term grassland biodiversity experiment with plant communities with either a history of co-occurrence (selected communities) or no such history (naïve communities) over a four-year period in which a major flood disturbance occurred.

We found selected communities had temporally more stable biomass than the same communities of naïve plants, especially at low species richness. Furthermore, selected communities showed greater short-term biomass recovery after flooding, resulting in more stable post-flood productivity. In contrast to a previous study, the positive diversity–stability relationship was maintained after the flooding. Our results were consistent across three soil treatments simulating the presence or absence of co-selected microbial communities. We suggest that prolonged exposure of plant populations to a particular community context and abiotic site conditions can increase ecosystem temporal stability and resistance to disturbance. We argue that selection during the course of a biodiversity experiment is the most parsimonious explanation for these effects. A history of co-occurrence can in part compensate for species loss, as can high plant diversity in part compensate for the missing opportunity of such adaptive adjustments.

## INTRODUCTION

Biodiversity experiments simulating the loss of plant species from grassland communities have shown that less diverse communities have reduced mean (Balvanera et al. 2006, Cardinale et al. 2012) and increased variation in aboveground biomass through time (Tilman et al. 1998, 2006, Hector et al. 2010). However, it is not clear whether these communities may regain functioning and stability over time while still being at low diversity. The few biodiversity experiments that lasted more than 10 years showed that functioning tended to decrease in low-diversity communities and to increase in high-diversity communities, leading to an increased slope of the biodiversity–biomass production relationship over time (Reich et al. 2012, Meyer et al. 2016, Guerrero-Ramírez et al. 2017). In one of these experiments, the Jena Experiment in Germany (Weisser et al. 2017), it was shown that divergent evolutionary changes of plant species in monocultures vs. mixtures during the first 8 years contributed to this strengthening of the biodiversity–functioning relationship (Zuppinger-Dingley et al. 2014, van Moorsel et al. 2018, 2019). Feedbacks between plants and soil organisms, however, had less explanatory power (van Moorsel et al. 2018, Schmid et al. 2019, Hahl et al. 2020).

Ecosystem resistance, recovery, and resilience that underlie stability may depend on plant diversity (Pfisterer and Schmid 2002, Isbell et al. 2015, Fischer et al. 2016). The mechanisms by which diversity stabilizes ecosystem biomass production are based on differences among genotypes or species in their responses to the abiotic or biotic environment (Schmid 1994, Tilman et al. 1998, Hector et al. 2010). For instance, different plant species may exhibit high performance under different environmental conditions (termed response diversity, Elmqvist et al. 2003, Isbell et al. 2011). Asynchrony among species performances in terms of biomass production, derived from interspecific differences in responses to environmental variation, can thus allow diverse communities to resist disturbance or recover to maintain performance, often referred to as insurance or portfolio effect (Yachi and Loreau 1999, Hector et al. 2010, Thibaut and Connolly 2013, de Mazancourt et al. 2013).

The development of stability over time in long-term biodiversity experiments has not been analyzed so far, but data from the Jena Experiment (Weigelt et al. 2010, Weisser et al. 2017) show that combined intra- and inter-annual biomass variation in experimental communities decreased over time for the first 8 years (i.e. ecosystem stability, measured as the inverse of the coefficient of variation of plant biomass, increased over time; Appendix S1: Fig. S1A), stability thus increased. During this time period, climatic variability did not decrease (Appendix S1: Fig. S1B, C, D), yet over time, as the stability of community biomass decreased, so did mean stability of interannual precipitation (inverse of variability). This correlation between climatic and biomass stability demonstrates a fundamental problem of interpretation in studies that confound (community) age and physical time, leading us to design an experiment that allows a separation of the two. Our overall hypothesis is that the increase in biomass stability over time in the Jena Experiment can at least in part be attributed to community age. As communities develop following sowing, species abundance distributions and gene frequencies over the years change, and such adjustments between and within species may increase stability (Strauss et al. 2006, Aubree et al. 2020). Comparing such “old” communities with “new” communities of the same species composition at the same time and under the same environmental conditions addresses our overall hypothesis as described below. Here we use “co-occurrence history” when we refer to plant species with a history of growing in the company of one another over a certain period of time, potentially developing stronger interactions or associations with both the plant and soil community partners over time.

Evidence that interactions between co-occurring species can become more stable has been obtained when long time spans were considered. For example, a prolonged period of co-occurrence increased “stabilizing differences” between phylogenetically distinct annual plant species in comparison with similar pairs of species without co-occurrence history (Germain et al. 2016). Here we ask if prolonged co-occurrence within a local community might already result in changed species interactions and reduced competition in the short term, such as during the course of a biodiversity experiment, and for perennial species. To exclude confounding effects of individual plant age and site conditions, we started our experiment with seedlings planted in equal species compositions and under equal soil conditions at the field site in Jena. In addition, we applied different soil treatments to assess the potential contribution of soil organisms that over time associate with plant communities and may (de)stabilize plant communities by changing nutrient provision and the plant’s health (Eisenhauer et al. 2011, 2012). Communities ranged in species richness from one, two and four to eight plant species. We refer to “old” communities as “*selected* communities” since they were assembled with offspring from individuals that had co-occurred in the same plots of the Jena Experiment over 8 years. We refer to “new” communities as “*naïve* communities” because they were assembled with offspring from seeds that were obtained from the original seed supplier for the Jena Experiment.

We partitioned our overall hypothesis into three more specific ones. (1) We previously found that selected communities were more productive than naïve communities in the same experiment at the 2- and 4-species richness levels but not at the 8-species richness level. We thus hypothesized that selected communities have more stable biomass than naïve communities and that differences in stability between selected and naïve communities are most pronounced at intermediate diversity. (2) Stability is further increased when plants are growing with their native soil organisms. These two hypotheses were assessed by analyzing the combined intra- and inter-annual variation in plant community biomass across seven seasonal aboveground harvests from 2012–2015 (Appendix S1: Fig. S2). A natural flood event (Blöschl et al. 2013) after the third harvest gave us the opportunity to additionally analyze the resistance, recovery, and resilience (Ruijven and Berendse 2010, Lloret et al. 2011, Hillebrand et al. 2018) of test communities in response to the flood and compare the pre- with the post-flood biomass stability using the first three and the last three harvests, respectively. Together, resistance and recovery determine ecosystem resilience as we define it here, namely how ecosystem biomass production differs between pre- and post-disturbance states (Lloret et al. 2011). Assuming that co-occurrence history should also increase stability towards perturbation, our third hypothesis is that (3) selected communities show greater resistance, resilience, and recovery in response to the flood event, thus addressing specifically the influence of the natural flood event as environmental stress.

## METHODS

### Field site

This study was conducted at the Jena Experiment field site (Jena, Thuringia, Germany, 51 °N, 11 °E, 135 m a.s.l.) from 2011–2015. The Jena Experiment is a long-term biodiversity field experiment located on the banks of the Saale River. In 78 experimental field plots of different diversity levels, 60 mostly perennial species typically forming species-rich grassland ecosystems under low-intensity management are grown in a number of species combinations since 2002 (Roscher et al. 2004).

### Co-occurrence (selection) history

This study included eleven monocultures, twelve 2-species mixtures, twelve 4-species mixtures and twelve 8-species mixtures for a total of 47 species compositions assembled from a pool of 49 species in the large plots of the Jena Experiment (Roscher et al. 2004). This subset of large plots excluded 16- and 60-species mixtures as well as monocultures and mixtures with very poor growth of some species to obtain nearly equal replication of communities at each diversity level (one initially chosen monoculture could not be used because it contained individuals of a different species from the one originally planted) (van Moorsel et al. 2018). The 49 species were mostly perennials and capable of outcrossing and represented the functional groups grasses (including graminoids of families other than Poaceae; 16 species), legumes (Fabaceae; 12 species) and herbs (21 species, Appendix S1: Table S5) (Roscher et al. 2004).

We used two co-occurrence-history treatments: communities assembled with offspring of plants that had grown together for 8 years in the 47 large plots of the Jena Experiment (“selected” communities, Appendix S1: Table S5) and communities assembled with plants without a common history of co-occurrence in the Jena Experiment (“naïve” communities). The naïve communities were naïve in the way that they had not experienced selection in communities in the Jena Experiment but have been exposed to selection in their original field sites and in the monoculture gardens of the seed supplier.

In total, there were 219 selected populations from different diversity levels in the Jena Experiment for the 49 species. The plants of naïve communities were grown from seeds obtained in 2010 from the same commercial supplier (Rieger Hofmann GmbH, in Blaufelden-Raboldshausen, Germany) who provided the seeds used for the establishment of the Jena Experiment in 2002. The supplied seeds for both the original seed lots in 2002 and the new seed lots in 2010 originated from various field sites in Germany and had been cultivated by reseeding every year for up to five years in monoculture. We could not use seeds from the original lots for the naïve communities because there was not enough seed material left, some species had low germination rates and we were concerned that the long storage might have affected seed quality. Even though the new seed lots from 2010 likely contained other genotypes than the original seeds lots from 2002, we here focused on the species- and community-level replication to test our evolutionary hypotheses. Therefore, we assume a random variation for potential biases between seed lots from 2002 and 2010 for each of the 49 species and for each of the 141 assembled communities (47 species compositions × 3 soil treatments). These biases could have inflated the error terms used in the hypothesis tests of the mixed models described below and thus reduced observed effect sizes for the term co-occurrence history. However, by using seeds from the same supplier rather than entirely different seed sources of the same 49 species, we intended to reduce this bias as much as possible.

To reduce potential maternal carry-over effects from the field, seeds of selected communities were produced in an experimental garden in Zurich, Switzerland, from cuttings that had been made in the Jena Experiment in 2010. Cuttings from multiple individuals per species were planted in Zürich in the original species combination in plots fenced with plastic netting to minimize cross-pollination between the plots and surrounded by concrete walkways and frequently mowed lawns to avoid pollinations from outside plants. To allow pollinator access the plots in the experimental garden were left open at the top (Zuppinger-Dingley et al. 2014). In a subset of experimental communities, seed production in Zürich was not sufficient. In those cases, additional seeds were collected directly in the plots of the Jena Experiment (see Appendix S1: Table S6). The “selected” seeds were thus offspring of plant populations that had been sown in 2002 and grown until 2010 in plots of the Jena Experiment plus — for most of the seeds — one season in the experimental garden in Zurich in the same species composition.

To make sure selected and naïve plants had similar starting conditions and to reduce differential maternal carry-over effects between the two co-occurrence histories, we germinated all seeds and propagated the resulting seedlings in a glasshouse at the same time and under the same environmental conditions. In January 2011, the seeds were germinated in potting soil (BF4, De Baat; Holland) and in March 2011 the seedlings were transported to the Jena Experiment field site and transplanted into 2 × 2 m smaller plots within the original large plots (see Fig. 1). There were four 1 × 1 m quadrats separated by plastic frames with different soil treatments in each 2 × 2 m plot (see next section) and each quadrat was split into two 1 × 0.5 m halves (“half-quadrats/subplots”, not separated by plastic but by a cord fixed in the ground). We planted seedlings of selected communities into one half and seedlings of naïve communities into the other half of each quadrat in a hexagonal pattern at a density of 210 plants per m^2^ with a 6-cm distance between individuals. By planting seedlings instead of sowing seeds, we could ensure equal abundances and positions of species in the 141 pairs of 1 × 0.5 m subplots containing the 282 test communities of different co-occurrence history, species diversity, and soil treatments. After transplanting, the seedlings received water every second day for six weeks. The experimental set up is shown in Fig. 1.

**FIG. 1.**
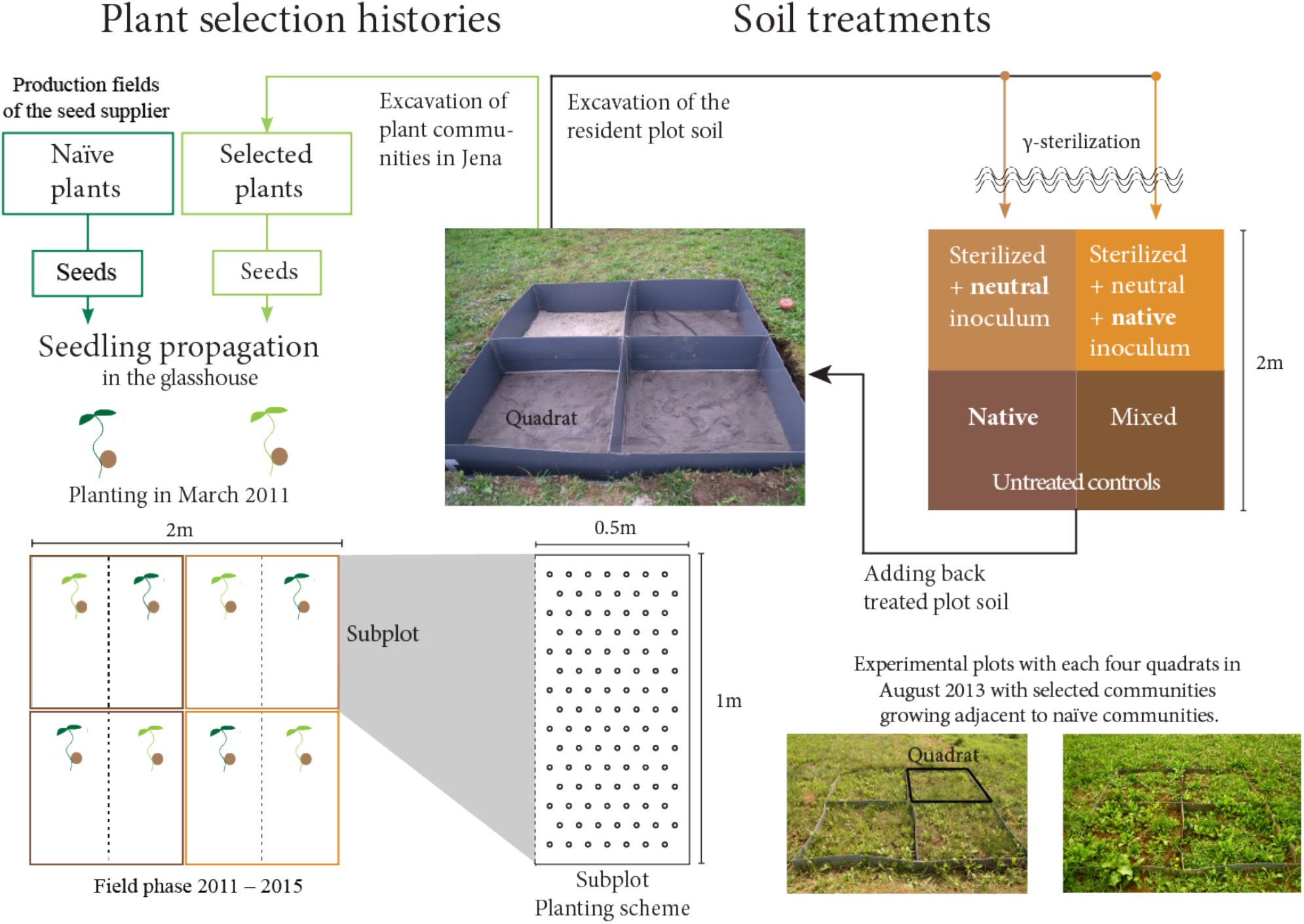
Experimental set-up of plant communities in the field. Seeds from plants that had been co-occurring for eight years in 47 plots of the Jena Experiment (selected plants) and seeds purchased from a seed supplier (naïve plants) were germinated at the same time in a glasshouse. These seedlings were then transplanted back to the Jena field site in March 2011 according to randomized planting schemes with equal species composition and abundances. Selected (light green) and of naïve communities (dark green) were grown, in the same 47 plots from which selected plants had been taken, in four quadrats separated by plastic frames with different soil treatments (unsterilized native or mixed soil or sterilized soil with native or neutral inoculum, see Methods). The mixed-soil treatment was not used in this paper because it was harvested early for a different experiment. Co-occurrence history (selected vs. naïve) was thus a split-split plot treatment replicated for 47 community compositions (including 11 monocultures) times three soil treatments. We ensured equal abundances and positions of species in the 141 pairs of 1 × 0.5 m subplots (see planting scheme).

### Soil treatments

Within each 2 × 2 m plot of the 47 large plots of the Jena Experiment, we removed the original plant cover in September 2010 and used it for the plant propagation in the experimental garden in Zurich (see previous section). Subsequently, we excavated the soil to a depth of 0.35 m, added a 10-cm layer of sand to the bottom of the plots and covered it with a 0.5-mm mesh net. We separated the borders of the plots and the quadrats by plastic frames. The excavated native soil from each of the plots was sieved and four soil treatments were prepared. Half of the soil (approximately 600 kg per plot) was γ-irradiated to remove the original soil biota. Half of the sterilized soil was then inoculated with 4% (by weight) of live sugar-beet soil and 4% of sterilized native soil of the corresponding plot (“neutral soil” obtained by inoculation). Live sugar-beet soil was added to create a neutral soil community and was previously collected in an agricultural sugar-beet field not associated with the Jena Experiment, but with comparable soil properties. The second half of the sterilized soil was inoculated with 4% (by weight) of live sugar-beet soil and 4% of live native soil of the corresponding plot (“native soil” obtained by inoculation). The non-sterilized part of the excavated soil was used for the second two soil treatments. Half of this soil was filled back into one quadrat of the corresponding plot (“native soil”). The other half of the unsterilized soil was mixed among all plots and filled into the remaining quadrats (“mixed soil”). However, this fourth soil treatment was destructively harvested for another experiment, which is why we excluded it from all analyses.

The soils were left to rest in closed bags to allow for the soil chemistry to equalize and to encourage soil biota of the inocula to colonize the sterilized soil before planting. The soils were then added into the quadrats in December 2010, and all quadrats were covered with a net and a water-permeable black sheet to avoid spilling between quadrats until seedling transplantation in March 2011.

In 2011 and 2012, we could show that the soil treatments were distinct one year and two years post establishment (van Moorsel et al. 2018). We sampled soil again at the end of the experiment in 2015 (2 years after the flood event) and found that soil treatments still differed strongly with regard to chemistry and microbial composition (Appendix S1: Table S4).

### Sampling of aboveground biomass

The plant communities were weeded three times a year and the plants were cut to 3 cm above ground twice a year at typical grassland harvest times (late May and August) in central Europe. Plant material from a 50 × 20 cm area in the center of each half-quadrat was collected to measure aboveground biomass. We sorted the biomass into species, dried it at 70°C and weighed the dried biomass. There were four May harvests (2012–2015) and three August harvests (2012–2014) because the experiment was terminated after the fourth May harvest in 2015.

### Natural flood event

In June 2013, the field site was flooded because of heavy rains in central Europe (Blöschl et al. 2013, Wright et al. 2015). The flood duration (maximum 25 days) and depth of water (maximum of 40 cm) varied between 2 × 2 m plots but not between co-occurrence-history and soil treatments within plots (Fischer et al. 2016). Because flood severity (Wright et al. 2015) did not influence the diversity–stability relationship or any other of the dependent variables in the present study (data not shown), we excluded flood severity indices from all analyses.

### Data analysis

#### Temporal stability of community biomass and climate

To address hypothesis 1, we first calculated the stability of community aboveground biomass as the inverse coefficient of combined intra- and inter-annual variation (*CV*_*com*_^−*1*^) among sequential spring and summer harvests. The stability of a single community was thus the mean community aboveground biomass (*μ*_*com*_) divided by its standard deviation (*σ*_*com*_). We deliberately combined seasonal and annual variation in the stability measure to encompass both aspects of temporal variation in a single measure. The basic sequence for this measure was spring year *n*, summer year *n*, and spring year *n*+1, which had shown increasing stability during the 8 selection years in the Jena Experiment (2003/4, 2005/6, 2007/8, 2009/10; see Appendix S1: Fig. S1A). This sequence allowed us to exclude the summer harvest 2013, which was taken shortly after the flood event and was used for the calculation of resistance and recovery (see below); and it increased the independence of the sequential measures from 2003–2010. We calculated interannual mean spring precipitation and temperature stability (Knapp 2001) for the same time intervals in Jena (see Appendix S1: Fig. S1B). Temperature and precipitation were measured with a weather station on site (see Appendix S1: Fig. S1C, D).

We also analyzed pre-flood (first three harvests) and post-flood (last three harvests) stability in the same way as described at the beginning of this section. Furthermore, we calculated the species compositional turnover between pre- and post-flood conditions. Because it includes species abundances, we used the Bray-Curtis dissimilarity between pre-flood (averaged over the first three harvests) and post-flood abundances of species (averaged over the last three harvests). Although the separate analyses of pre- and post-flood stabilities are partly confounded with the analysis of overall stability across the three pre- and three post-flood harvests (see previous paragraph) we did both types of analyses to focus on different aspects of stability. Whereas the analysis of the overall stability as an integrative measure allowed us to better estimate contributions of asynchrony and population stability to community stability (see next section), the separate analyses of pre- and post-flood stabilities allowed us to test specifically if the flooding event not only affected resistance, recovery, and resilience of communities (see below), but also the temporal stability over time in absence of further perturbations.

#### Population stability and species asynchrony

We calculated average stability of biomass at the population level (*CV*_*pop*_^−*1*^) and community-wise species biomass asynchrony (1−*θ*) over the same time span as overall stability. Stability of biomass at the population level was calculated as the average stability of biomass of individual species following Thibaut and Connolly (2013). Asynchrony was calculated as the “synchrony index” (*θ*) following Loreau and de Mazancourt (2008), which ranges between 0 and 1, thus, asynchrony is 1-*θ*. For monocultures, population stability equals community stability and asynchrony is zero (*θ* is 1). Community stability is the product of population stability and the square root of species synchrony (Thibaut and Connolly 2013, de Mazancourt et al. 2013). This allowed us to assess these two components of community stability separately.

#### Resistance, recovery, and resilience

To address hypothesis 3, we calculated resistance, recovery, and resilience measures (Schläpfer and Schmid 1999, Ruijven and Berendse 2010, Hillebrand et al. 2018) in response to the natural flood event in 2013 (see Fig. 3). Resistance is the difference in community biomass between the average of the three harvests before the flood and the community biomass directly after the flood (summer 2013), more negative values indicating lower resistance. Recovery is the difference between the biomass produced after recovery from the flood (averaged over the three last harvests) and the biomass directly after the flood (summer 2013), where positive values indicate the amount of biomass recovered. Resilience is the difference between the average biomass of the three harvests before the flood and the average biomass of the three harvests after recovery. Values close to zero or positive values indicate that communities had returned or overshot their pre-flood state, respectively, after the flood; and negative values indicate that post-flood biomass had not returned to its pre-flood state.

#### Statistical analysis

Variation in community stability, synchrony, and population stability was analyzed with linear mixed-effects models. Stability measures were log-transformed to improve homoscedasticity and obtain normally distributed residuals in the analyses (Schmid et al. 2017). Fixed-effects terms were plant species richness (log scale, addressing hypothesis 1), co-occurrence history (selected vs. naïve communities, addressing hypothesis 1), and soil treatment (native, inoculated-native or inoculated-neutral soil, addressing hypothesis 2). Plots and quadrats were used as random-effects terms to get appropriate errors for significance tests (Schmid et al. 2017). We added all significant interactions of the fixed-effects terms as additional fixed-effects terms to the models (see Table 1). For reasons of consistency and to allow the use of all data in analyses with covariates, we included monocultures in the analysis of asynchrony. For graphical displays of relationships between species richness and stability measures and asynchrony, means across soil treatments were corrected for differences between plots within species-richness levels, which corresponds to using plots and quadrats in the mixed-model analyses. Because co-occurrence history was a split-plot/split-quadrat treatment applied within each quadrat, it was not affected by the correction. The corrections were obtained by fitting a model with plots and quadrats only and adding the residuals to the diversity-level means. This correction corresponds, for example, to that used in metabolic ecology, when metabolic rate corrected for temperature is plotted against body size (Brown et al. 2004).

**TABLE 1.** Mixed-model ANOVA results for log-transformed community stability, log-transformed mean population stability and untransformed asynchrony.

| | Stability ($CV^{-1}$) | | | | Population stability ($CV_{pop}^{-1}$) | | | Asynchrony ($1-\theta$) | | |
| --- | --- | --- | --- | --- | --- | --- | --- | --- | --- | --- |
| | $DF_n$ | | | | | | | | | |
| Fixed terms | $um$ | $DF_{den}$ | $F$ | $P$ | $DF_{den}$ | $F$ | $P$ | $DF_{den}$ | $F$ | $P$ |
| Log richness ($R_{log}$) | 1 | **44.1** | **10.74** | **0.002** | **44.1** | **5.27** | **0.027** | **44.1** | **143.00** | **<0.001** |
| Soil treatment (SH) | 2 | 87.1 | 0.64 | 0.529 | 87.1 | 1.30 | 0.278 | 87.1 | 0.87 | 0.424 |
| Co-occurrence history (CH) | 1 | 135.0 | 1.80 | 0.181 | 135.0 | 3.79 | 0.054 | 135.0 | 0.50 | 0.479 |
| SH x $R_{log}$ | 2 | 87.2 | 0.05 | 0.954 | 87.2 | 0.01 | 0.992 | 87.2 | 0.38 | 0.685 |
| CH x $R_{log}$ | 1 | **135.0** | **4.79** | **0.030** | **135.0** | **8.38** | **0.004** | 135.0 | 0.05 | 0.830 |
| Random terms | $N$ | $Var. 10^{-3}$ | $SE 10^{-3}$ | | $Var. 10^{-3}$ | $SE 10^{-3}$ | | $Var. 10^{-3}$ | $SE 10^{-3}$ | |
| Plot | 46 | 100.1 | 25.9 |  | 95.6 | 23.5 |  | 17.9 | 4.6 |  |
| Plot x SH | 137 | 15.5 | 10.9 |  | 13.9 | 7.4 |  | -0.1 | 2.0 |  |
| Residual | 274 | 92.3 | 11.2 |  | 58.4 | 7.1 |  | 20.0 | 2.5 |  |

*Notes:* The effects of species richness (log scale), soil treatments, and co-occurrence history on the stability of community and population biomass and on asynchrony across the entire experimental period from 2012–2015 were analyzed (excluding the time point immediately after the extreme event of a late spring flood in June 2013). Bold italic text highlights significant effects. $DF_{num}$ = numerator degrees of freedom, $DF_{den}$ = denominator degrees of freedom, $F$ = variance ratio, $P$ = probability of type-I error.

Variation in resistance, recovery, and resilience was also analyzed with the same linear mixed-effects models as described above. Since the measures of resistance, recovery, and resilience can depend on the magnitude of the pre-flood biomass (Pfisterer and Schmid 2002, Wright et al. 2015), we analyzed additional models, which included the average of the three harvests before the flood as covariate (see Appendix S1: Table S1).

To compare the magnitude of the plant community response to either biodiversity or co-occurrence, we calculated effect sizes. Because our experimental design was nearly orthogonal, general linear models could replace the linear mixed models to obtain the sum of squares (SS) for all terms (Schmid et al. 2017). We calculated the total SS for all fixed-effects terms as 100% and then used the %SS of each individual fixed-effects term as effect size, which allowed us to compare the relative explanatory power of the different fixed-effects terms (Grömping 2006) (see Appendix S1: Fig. S3).

All analyses were conducted using the software R, version 3.2.4 (R Development Core Team 2017). Mixed models using residual maximum likelihood (REML) were fitted using the package ASReml for R (Butler 2009) and the package ‘Pascal’ available at GitHub (Schmid et al. 2017).

## RESULTS

### Co-occurrence history partially compensates the negative effects of biodiversity loss on biomass stability

Community biomass stability across pre-flood and post-flood harvests increased with species richness (Figure 2A, Table 1). Differences in community biomass stability between soil treatments were insignificant (Table1). Differences between selected and naïve communities (co-occurrence treatment) were small, however, at low diversity, selected communities were more stable than naïve communities, reflected by a significant co-occurrence history x species richness interaction (Table 1; Fig. 2A).

**FIG. 2.**
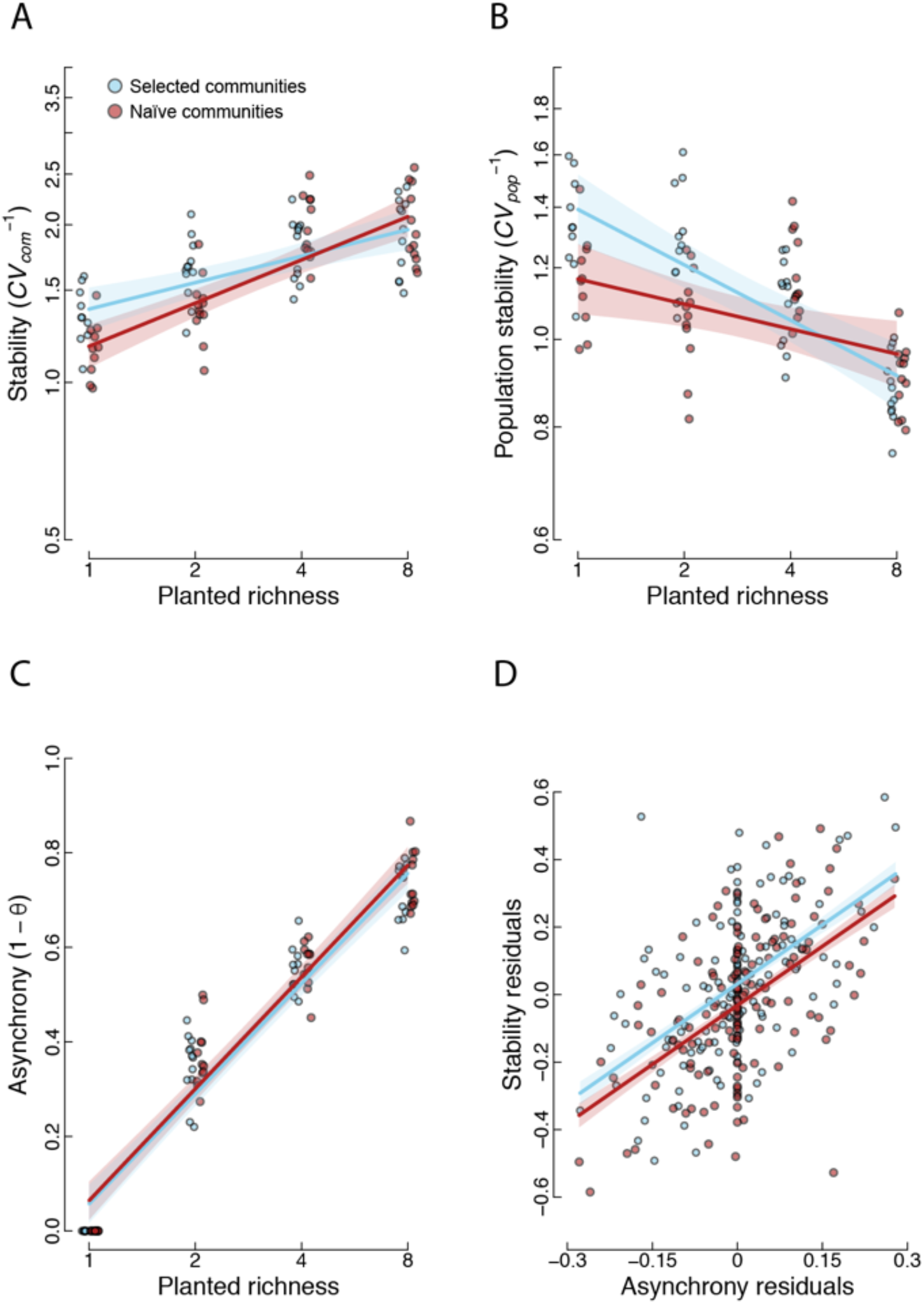
The biodiversity–stability relationship for selected (blue) and naïve communities (red). **(A)** Community stability, (**B)** mean population stability, **(C)** asynchrony, **(D)** relationship between stability and asynchrony after correction for all other model terms except co-occurrence history. The corrections were obtained by fitting a model with plots and quadrats only and adding the residuals to the diversity-level means (see Methods). Colored bands show standard errors of predictions from mixed models as presented in Table 1. For significances see Table 1 (panels A–C); the slopes in panel D are significant at *P* < 0.001. In panels A–C points are means of the three soil treatments estimated from the model in Table 1. Points in D are residual values of each plant community after accounting for the variation due to soil treatments, planted richness, and plot identity.

Population biomass stability decreased with species richness, but at low diversity, the population biomass stability was also higher in selected communities (Table 1; Fig. 2B). In contrast, species asynchrony in terms of biomass increased for both selected and naïve communities in parallel with increasing species richness (Table 1; Fig. 2C). When we corrected community stability and species asynchrony for all model terms except co-occurrence history (i.e. taking residuals after fitting the plot x soil treatment interaction), stability residuals strongly increased with asynchrony residuals (P < 0.001), and selected communities, in addition, were consistently more stable than naïve communities (P < 0.01; Fig. 2D).

An analysis of effect sizes showed that log-transformed richness had the strongest effect (between 77 and 99%, Appendix S1: Fig. S3A) on community stability, population stability, and asynchrony.

### Diverse communities were less resistant to a flood event but recovered better

A naturally occurring flood in early summer 2013 strongly reduced biomass in that summer (Fig. 3 and Appendix S1: Fig. S2). However, in contrast to the main plots in the Jena Experiment (Wright et al. 2015), the flood did not interfere with the positive diversity–community biomass stability relationship in our smaller plots (Fig. 5). In the short term, diverse communities, especially selected ones, were the least resistant (Fig. 4A). At low diversity, selected communities tended to have higher resistance than naïve communities, especially when adjusting for community biomass before the flood (by adding pre-flood biomass as a term in the model, see Appendix S1: Table S1; Fig. S4A).

**FIG. 3.**
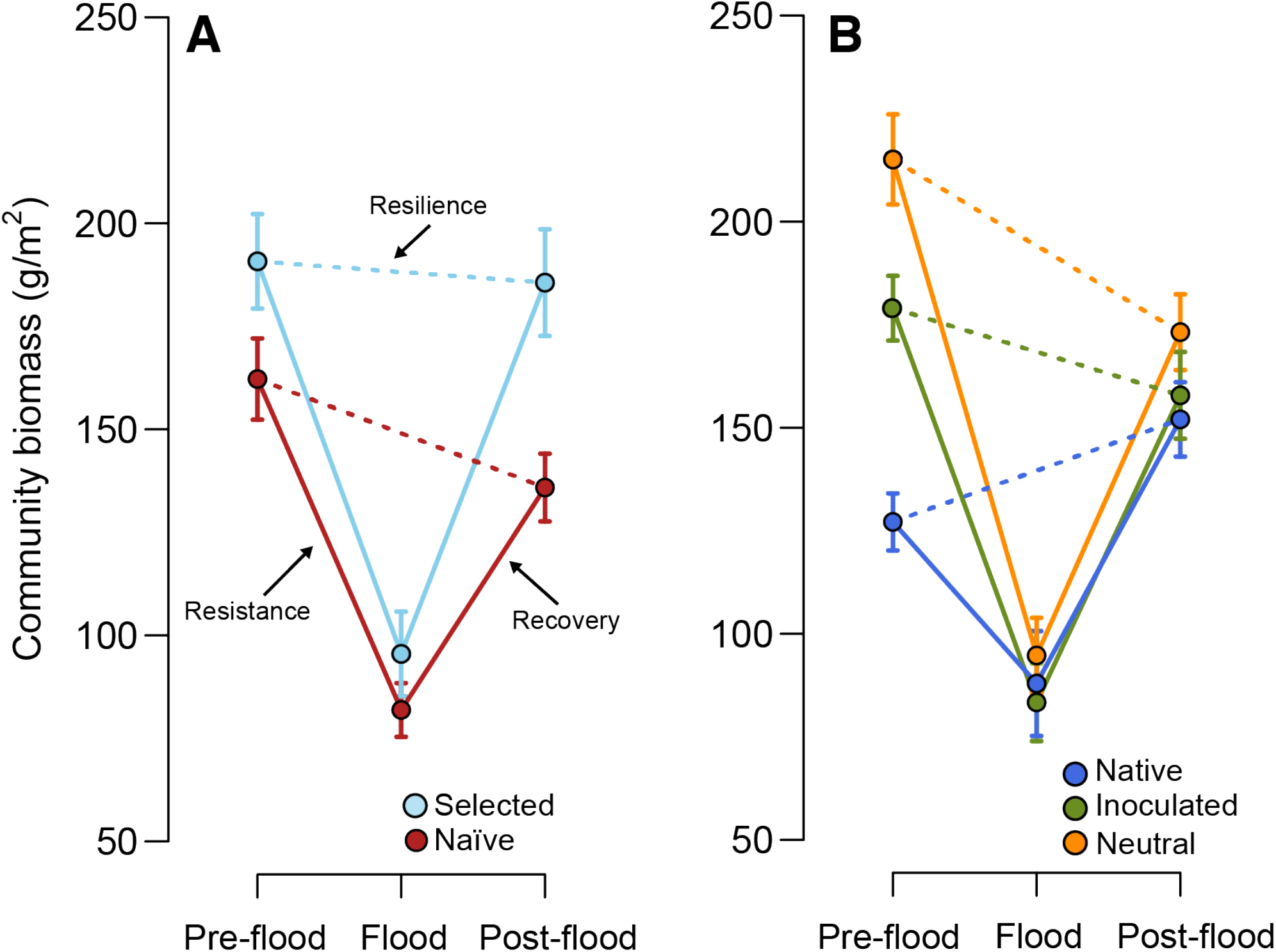
Plant community biomass before and after the flood event. Points indicate the average community biomass across all diversity levels for **(A)** selected (blue) and naïve communities (red) and **(B)** native soil (blue), sterilized soil with native inoculum (“inoculated”, green) and sterilized soil with neutral inoculum (“neutral”, orange). Resistance is the difference in biomass between the average of the three harvests before the flood (May 2012, August 2012, and May 2013) and the biomass directly after the flood (label “Flood” on x-axis corresponding to summer harvest in August 2013). Recovery is the difference in biomass between the average of the three harvests after recovery from the flood (May 2014, August 2014, and May 2015) and the biomass directly after the flood (“Flood” label). Resilience is the difference in biomass between the average of the three harvests after recovery from the flood and the average of the three harvests before the flood. See also Figure Appendix S1: S2. Means and standard errors were calculated from raw data.

**FIG. 4.**
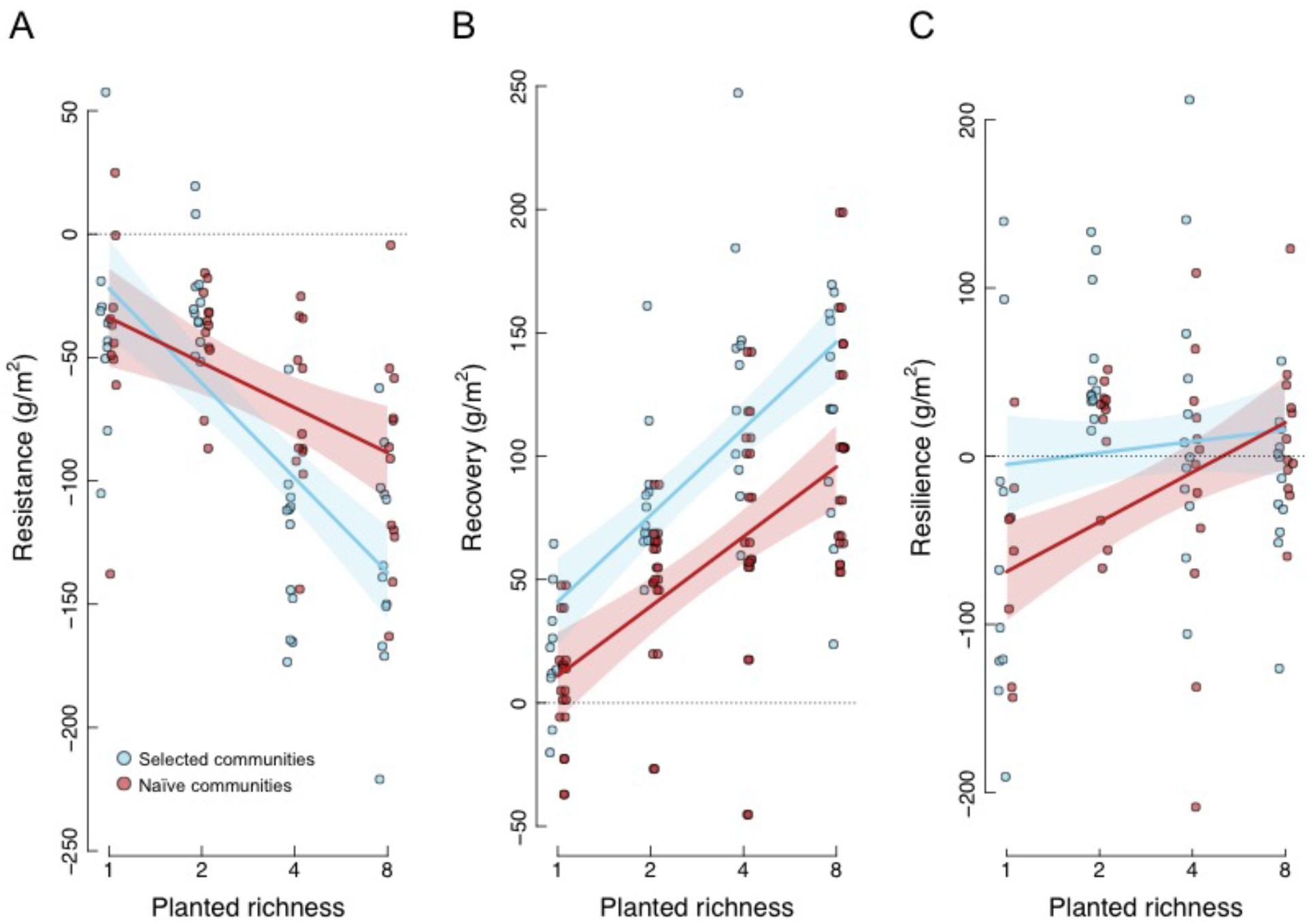
Resistance, recovery, and resilience to the flood event. **(A)** Biodiversity–resistance relationships, **(B)** biodiversity–recovery relationships, and **(C)** biodiversity–resilience relationships for selected (blue) and naïve communities (red). Colored bands show standard errors of predictions from mixed models as presented in Table 2. For significances see Table 2. Points are means of the three soil treatments estimated from the model in Table 2. The dashed line at 0 indicates no change in biomass in response to the flood (resistance), after the flood (resistance), or between pre- and post-flood harvests (resilience). Similar plots with values corrected for variation in pre-flood biomass as covariate are shown in Appendix S1: Fig. S3.

**FIG. 5.**
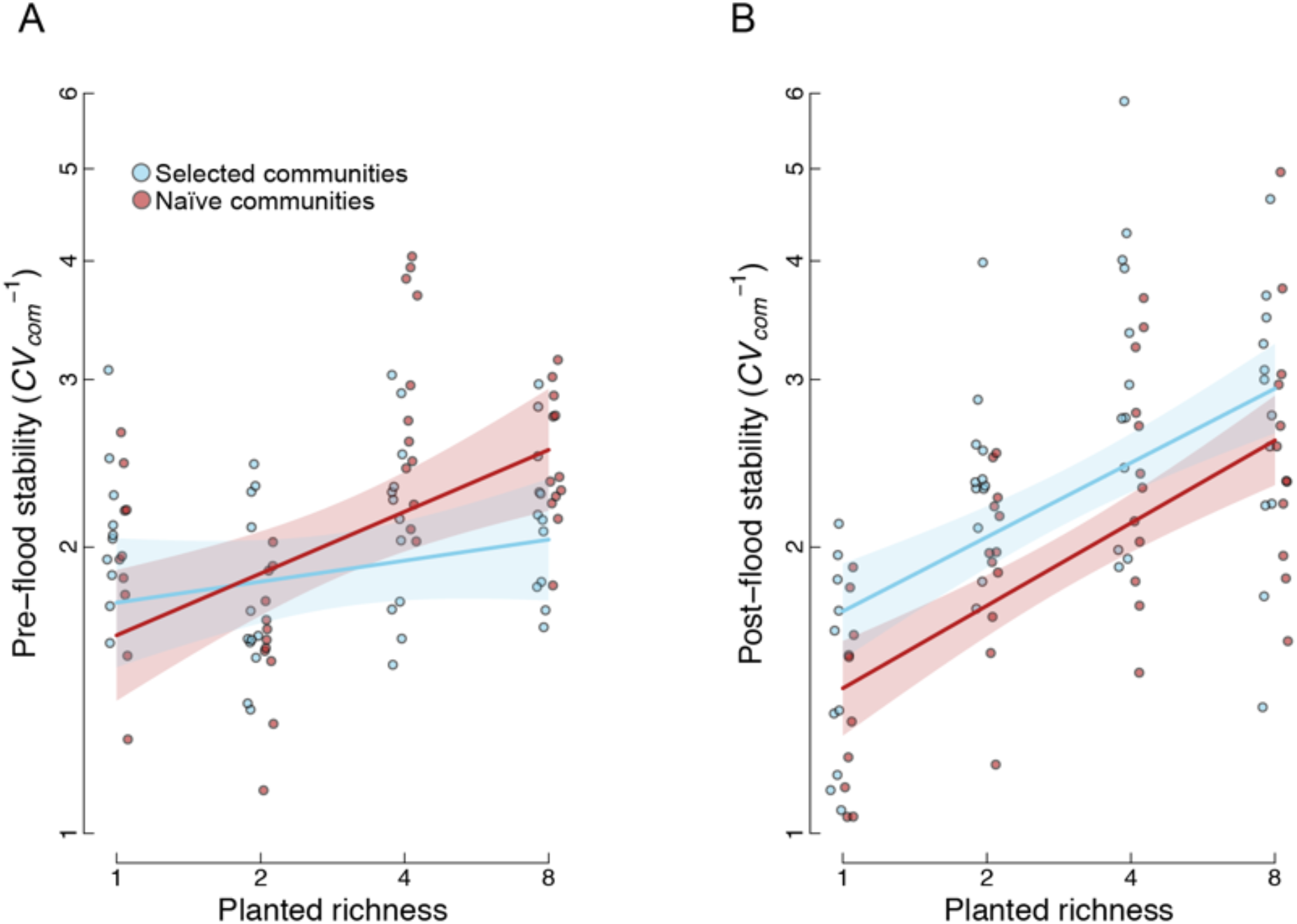
The biodiversity–stability relationship for selected (blue) and naïve communities (red). **(A)** The three harvests before the extreme event of a late spring flood in June 2013 and **(B)** the three harvests after recovery from the flood. Colored bands show standard errors of predictions from mixed models as presented in Appendix S1: Table S2. *P* < 0.001 for the effect of log richness in post-flood stability and *P* = 0.027 for the effect of co-occurrence history on post-flood stability. For other test-statistics see Appendix S1: Table S2. Points are means of the three soil treatments estimated from the model in Appendix S1: Table S2.

Plant communities in the non-sterilized native soil had the lowest biomass prior to the flood, lost the smallest amount that summer, and were thus most resistant (Fig. 3B). In contrast, plant communities grown in neutral soil had the highest biomass prior to the flood and were the least resistant to the flood resulting in a significant effect of soil treatment on resistance (Table 2; Fig. 3B). However, after first accounting for the pre-flood biomass, there were no effects of soil treatments on resistance (Appendix S1: Table S1).

**TABLE 2.** Mixed-model ANOVA results for resistance, recovery, and resilience of community biomass in response to the extreme event of a late spring flood in June 2013.

|  | Resistance |  |  |  | Recovery |  |  | Resilience |  |  |
| --- | --- | --- | --- | --- | --- | --- | --- | --- | --- | --- |
| Fixed terms | *DF<sub>num</sub>* | *DF<sub>den</sub>* | *F* | *P* | *DF<sub>den</sub>* | *F* | *P* | *DF<sub>den</sub>* | *F* | *P* |
| Log richness ($R_{log}$) | 1 | **44.2** | **9.41** | **0.004** | **44.1** | **15.95** | **<0.001** | 44.2 | 1.69 | 0.200 |
| Soil treatment (SH) | 2 | **87.3** | **14.07** | **<0.001** | 87.2 | 0.29 | 0.745 | **87.3** | **6.12** | **0.003** |
| Co-occurrence history (CH) | 1 | **135** | **4.19** | **0.043** | **135** | **14.50** | **<0.001** | 135 | 3.48 | 0.064 |
| SH x $R_{log}$ | 2 | **87.5** | **5.95** | **0.004** | 87.4 | 1.73 | 0.184 | **87.5** | **6.97** | **0.002** |
| CH x $R_{log}$ | 1 | **135** | **5.32** | **0.023** | 135 | 0.48 | 0.488 | 135 | 2.65 | 0.106 |
| Random terms | *N* | *Var.* | *SE* |  | *Var.* | *SE* |  | *Var.* | *SE* |  |
| Plot | 46 | 3645 | 1074 |  | 2234 | 771 |  | 6910 | 2238 |  |
| Plot x SH | 137 | 775 | 702 |  | -158 | 745 |  | 1933 | 1784 |  |
| Residual | 274 | 6246 | 760 |  | 7851 | 956 |  | 15914 | 1937 |  |

*Notes:* The effects of species richness (log scale), soil treatments, and co-occurrence history on responses of community biomass to flooding were analyzed. Bold italic text highlights significant effects. (Similar ANOVAs with pre-flood biomass as covariate are shown in Appendix S1: Table S3.) $DF_{num}$ = numerator degrees of freedom, $DF_{den}$ = denominator degrees of freedom, $F$ = variance ratio, $P$ = probability of type-I error.

Recovery of community biomass after the flood increased with species richness and was higher in selected than in naïve communities across all diversity levels and soil treatments (Table 2 and Appendix S1: Table S1; Fig. 4B and Appendix S1: Fig. S2B). Selected communities were also more resilient than naïve communities, as shown by the comparison of community biomass before and after recovery (Fig. 3A), and particularly at low diversity (Fig. 4C). However, the effect of co-occurrence on resilience was only significant if adjusted for pre-flood community biomass (Table 2 and Appendix S1: Table S1; Fig. 4C and Appendix S1: Fig. S4C). The three soil treatments strongly differed in their resilience, which averaged out their pre-flood differences in community biomass (Table 2; Fig. 3B).

Effect sizes (%SS) show that species richness had the strongest impact on resistance (42%), the interaction between soil history and species richness the strongest impact on recovery (37%) and species richness the strongest impact on resilience (70%, Appendix S1: Fig. S3B). Co-occurrence history contributed with 23% to resilience.

### Selected communities were more stable after the flood

We compared the combined intra- and interannual biomass stability over the first three harvests before the flood event (2012–2013, Fig. 5A) with the last three harvests after recovery (2014–2015, Fig. 5B). Before the flood, selected communities were not more stable than naïve communities, but there was a marginally significant interaction with species richness (Appendix S1: Table S2), which was similar in direction to the one of the overall community biomass stability across the entire experimental period. After the flood event, the selected communities were consistently more stable than the naïve communities across all diversity levels (Appendix S1: Table S2). Lastly, species turnover rates (Bray-Curtis similarity) between pre-and post-flood species compositions were not influenced by co-occurrence history or soil treatments, although they increased with species richness (Appendix S1: Table S3 and Fig. S5).

## DISCUSSION

Strengthening biodiversity–biomass production relationships observed in long-term biodiversity experiments (e.g. Reich et al. 2012, Guerrero-Ramírez et al. 2017) indicate a deterioration of monocultures and low-diversity mixtures and increase of functioning in more diverse mixtures (Meyer et al. 2016, Guerrero-Ramírez et al. 2017). We previously found that, in comparison with naïve communities, selected low-diversity mixtures were more productive than mixtures with higher diversity (van Moorsel et al. 2018). Here, we show that selected communities from the Jena Experiment also showed greater community biomass stability in comparison with naïve communities, particularly at low diversity.

### Selected communities recover better from an extreme event and are more stable at low diversity

Diverse communities are more stable in the face of disturbances (Isbell et al. 2015), such as a flood as happened to our test communities halfway through the experiment in June 2013 (Wright et al. 2015). Considering predicted future climate scenarios with increased frequency of extreme events (Stocker et al. 2013), including floods (Hirabayashi et al. 2013), this aspect of stability may even be more relevant than temporal stability under unperturbed conditions (Donohue et al. 2016). Further, both temporal stability and resistance to perturbations need to be considered because they may not always be positively correlated (Pennekamp et al. 2018).

In our experiment, diversity reduced ecosystem resistance in the short term, in line with previous findings for example in micro-ecosystems with ciliates responding to warming (Pennekamp et al. 2018). This was because 4- and 8-species communities had more biomass before the flood than 2-species mixtures and monocultures and thus could also lose more biomass (in absolute terms), a result found previously for community responses to drought (Pfisterer and Schmid 2002, Wang et al. 2007, Ruijven and Berendse 2010) and flood (Wright et al. 2015). Because selected communities were additionally more productive than naïve communities at the 8-species richness level, naïve communities were more resistant than selected communities as they had less to lose (see Fig. 4A). Diverse communities made up for their reduced resistance by increased recovery, as often found in biodiversity experiments (Ruijven and Berendse 2010, Lloret et al. 2011, but see Isbell et al. 2015). Remarkably, however, selected communities showed greater recovery than naïve communities along the entire species-richness gradient. In combination, the differential responses regarding resistance and recovery caused selected communities at low diversity to be more resilient than naïve communities, whereas no differences in resilience between selected and naïve communities were observed at higher diversity (see Fig. 4C). Some communities, mostly selected 2- and 4-species communities and both selected and naïve 8-species communities, were more productive after the flood than ever before (reflected in the positive resilience values shown in Fig. 4C). This could have been due to several potential non-exclusive causes: 1) continued accumulation of belowground biomass potentially less affected by flooding (and higher in selected than in naïve communities), 2) relative accumulation of beneficial microbes in comparison to plant antagonistic microbes (especially in sterilized soil treatments), 3) resource enrichment associated with the flood (as suggested by Wright et al. 2015). Accumulation of beneficial soil microbes seems to play a minor role though because in our analyses the interaction between soil treatments and co-occurrence history was not significant; that is, soil treatments did not differentially affect selected vs. naïve communities. But it is conceivable that selected communities could benefit more from resource enrichment because they had evolved better division of labor (Zuppinger-Dingley et al. 2014).

Whereas the differences in community temporal biomass stability between selected and naïve communities were only positive in monocultures before the flood (see Fig. 5A), the selected communities showed increased post-flood stability at all diversity levels (see Fig. 5B and Appendix S1: Table S2). This was driven by the improved recovery of the selected communities which resulted in a larger increase in mean biomass (van Moorsel et al. 2018) than in temporal variation of biomass and a consequently reduced CV of biomass. This improved stability of selected monocultures and mixtures after the flood event was likely due to local adaptation of plants to the abiotic conditions at the Jena field site, which is a natural floodplain, where the plant communities were exposed to soil moisture saturation at previous milder flood events in winter 2003 and winter 2005 (personal communication with C. Roscher). Selection may thus have favored individuals with traits that allowed them to perform well under such conditions and to recover more rapidly (Garssen et al. 2015, Wright et al. 2017). The contribution of such parallel evolutionary responses among the multiple species of our experiment to their abiotic environment was reflected in their increased population stability at low diversity (see Fig. 2B) and the consistently higher stability of selected communities across the entire range of species asynchronies (see Fig. 2D). However, in mixtures, adaptation to the biotic environment, i.e. species interactions, must also have been involved because the differences between selected and naïve communities depended on diversity. Because we did not detect any altered species abundance distributions (Vogel et al. 2019), it seems likely that changes in genotype frequencies within species, i.e. evolution in the community context (Strauss et al. 2006), contributed to increased stability. Genetic analyses on a subset of five herbaceous species from the Jena Experiment confirmed for one annual species and two perennial species the potential for such rapid evolutionary changes and their genetic basis, with consequential epigenetic and phenotypic changes (van Moorsel et al. 2019). Furthermore, we found quantitative-genetic divergence in eleven species, even though for those we did not do molecular analyses (Zuppinger-Dingley et al. 2014). The changes in genotype frequencies within species in selected communities could be attributable to differential mortality, growth, or reproduction among the initially sown genotypes (Barrett and Schluter 2008), recombination during sexual reproduction or, least likely, to mutation.

### Diverse communities recovered better regardless of co-occurrence history

At the highest diversity level, differences between selected and naïve communities were small, which was also reflected by their similar resilience (Fig. 4C). Thus, the difference between selected and naïve communities at the 8-species level was only (but still) visible in the larger dip that the selected communities took (i.e. the more negative resistance and the more positive recovery). The small differences between selected and naïve communities at the 8-species richness level mirror earlier findings for productivity, where mean yearly biomass was similar for selected and naïve communities at the 8-species richness level (van Moorsel et al. 2018). Potential effects of co-evolution may be weaker at higher diversity with less consistent and stable interactions between particular species (Connell 1980, van Moorsel et al. 2018). Stronger selective pressure between particular species leading to co-evolution could explain why the differences between selected and naïve communities were stronger at lower diversity, especially in 2- and 4-species mixtures. The increased resilience of selected communities at the lower diversity levels may in part also have been driven by evolutionarily increased facilitation (Bronstein 2009), which has been demonstrated for these plants in the Jena Experiment (Schöb et al. 2018). This would be in line with predictions that environmental stress might select for more positive interactions between species in plant communities (Callaway et al. 2002).

Resilience was slightly overshooting at the 8-species richness level (Fig. 4C), which indicates that species richness *per se* is already beneficial in the way that at lower richness, communities, in general, did not recover back to pre-flood biomass, i.e. they were not fully resilient. The increased resilience in selected and naïve 8-species communities was driven by a high recovery that overshot pre-flood levels of biomass production. The recovery of both selected and naïve communities may have been aided by the same causes as those mentioned in the previous section, namely higher belowground biomass or greater resource enrichment in more diverse communities. However, in contrast to Wright et al. (2015), we found that flooding did not decrease community stability and that after flooding diverse communities were still more stable than less diverse communities. Some of these dissimilarities might have been due to differences in the calculation of stability measures, in the range of species diversities tested, and in the management of experimental plots between the two studies.

### Population stability decreased with increasing species richness

Temporal stability in terms of biomass at the community level in grassland ecosystems can be driven by asynchronous population dynamics of species, allowing high compensatory population variation to be combined with low community-level variation over time (Flynn et al. 2008, Isbell et al. 2009, Hector et al. 2010, de Mazancourt et al. 2013, Gross et al. 2014). As shown before (e.g. Tilman et al. 2006), we found that community biomass stability increased but population biomass stability decreased with increasing species richness. However, this effect of species richness on population stability was weaker in naïve communities (see Fig. 2B), suggesting that adaptation to the abiotic environment partially compensated for the reduced species richness over time, especially in monocultures and low-diversity mixtures. In low-diversity mixtures, population stability could also have been increased due to reduced competitive interactions between plant species, consistent with the findings of evolutionary niche differentiation (Zuppinger-Dingley et al. 2014) and increased facilitation (Schöb et al. 2018) among species in mixtures in the Jena Experiment. By extension, similar evolutionary processes may have occurred between genotypes within monocultures, again consistent with previous findings showing evolutionarily changed phenotypic variation within monocultures after eight years of selection in the Jena Experiment (van Moorsel et al. 2018). The evolution of reduced inter- and intraspecific competition and parallel adaptations among the multiple species to the local abiotic conditions are mutually non-exclusive alternative explanations for the increased population stability at low diversity. Because community stability is the product of species stability and species synchrony (Thibaut & Connolly, 2013), yet asynchrony did not differ between selected and naïve communities (see Fig. 2C), we conclude that asynchrony did not contribute to the greater community stability of selected communities at low diversity.

### Influence of associated soil organisms

In our experiment we also wanted to investigate whether the association with native soil organisms over time might be a major factor in the observed strengthening of biodiversity–ecosystem functioning relationships in longer-term experiments (Reich et al. 2012, Meyer et al. 2016, Guerrero-Ramírez et al. 2017, this study). A number of studies showed that soil communities can strongly affect biodiversity effects in plant communities (Lau and Lennon 2012, van der Putten et al. 2013, terHorst et al. 2014). Specifically, for the Jena Experiment, previous findings suggested differential evolution of plant–soil feedbacks in monocultures vs. mixtures (Zuppinger-Dingley et al. 2016). We, therefore, deliberately designed our experiment with three soil treatments to detect possible effects of associated microbial communities on the stability of selected and naïve plant communities. However, although these soil treatments did produce significantly different microbial communities, we could not find any interactions between them and plant community co-occurrence history. Based on this “negative” result, we tentatively conclude that our above interpretations about plant evolutionary changes due to co-occurrence history were not confounded by differential assembly of soil communities over time in the Jena Experiment. That the soil treatments did work in principle could be seen by the main effects. Pre-flood productivity was lower when native soil biota were present, which could have been due to a greater density of antagonistic soil biota in native and native-inoculated soils (Schnitzer et al. 2011), or a greater pool of available soil resources resulting from the soil sterilization process in the two inoculated soils (Gebremikael et al. 2015). Recovery and resilience were higher for communities growing in native soil (see Fig. 3B), suggesting that native soil organisms did have a beneficial effect on both selected and naïve plant communities after they had been affected by the flood event.

## Conclusions

So far, evolutionary mechanisms underlying ecosystem stability in biodiversity experiments have only been studied in terms of phylogenetic relatedness that reflects evolutionary processes over long time scales, with conflicting results (e.g., Cadotte et al. 2012, Venail et al. 2015). Experimental evidence for short-term evolution leading to changes at the community level referred to as community evolution (van Moorsel et al. 2018), has been reported for microbial ecosystems (Gravel et al. 2011, Lawrence et al. 2012, Fiegna et al. 2014, 2015, Zhao et al. 2016). More recently, a modelling approach demonstrated the remarkable potential of co-adaptation to modify biodiversity–productivity relationships (Aubree et al. 2020). Short-term evolutionary processes could be particularly relevant in plant communities facing rapid global change (Schmid et al. 1996, Davis et al. 2005), including extreme climatic events; and for conservation and restoration approaches (Nagel et al. 2019), because plants are fixed in place and can only move by propagule dispersal. Here we have shown that evolutionary changes affecting biomass stability may already take place after few generations of sexual reproduction in communities of perennial plant species, likely due to sufficient “standing genetic variation” (Fakheran et al. 2010) in the original seed populations used at the start of the Jena Experiment (van Moorsel et al. 2019). We conclude by suggesting that others with long-term biodiversity experiments do similar follow-up experiments. Comparable results from a number of biodiversity experiments around the globe will strengthen the hypothesis that selection in a community context can increase stability, which would have far-reaching consequences for the fields of conservation and restoration ecology.

## Additional information

**Supplementary information** is available for this paper online.

## Data and code availability

Data and code (R scripts) are available from the corresponding author and will be made publicly available upon acceptance on the Pangaea repository.

## Acknowledgements

Thanks to Debra Zuppinger-Dingley, Dan Flynn, and Varuna Yadav for the establishment of the experimental plots. Thanks to the Jena Experiment for providing infrastructure and help and to D. Trujillo and M. Furler for technical assistance. This study was supported by the Swiss National Science Foundation (grant numbers 130720, 147092 and 166457 to B.S.) and the University Research Priority Program Global Change and Biodiversity of the University of Zurich. The Jena Experiment is funded by the Deutsche Forschungsgemeinschaft (DFG, German Research Foundation, FOR 1451) with additional support by the Friedrich Schiller University Jena and the Max Planck Institute for Biogeochemistry in Jena. NE acknowledges support from the German Centre for Integrative Biodiversity Research (iDiv) Halle-Jena-Leipzig, funded by the German Research Foundation (DFG FZT 118).

## Author contributions

B.S. designed research, T.H. and S.J.V.M. performed research; A.E. and N.E. maintained and coordinated the field site; S.J.V.M., C.W. and B.S. analyzed data; S.J.V.M., B.S., O.L.P. and C.W. wrote the paper. All co-authors contributed to subsequent versions of the paper.

