## Supplemental Information for "Co-occurrence history increases ecosystem temporal stability and recovery from a flood in experimental plant communities"

**Ecology**

**Appendix S1 for:**

### Co-occurrence history increases ecosystem temporal stability and recovery from a flood in experimental plant communities

Sofia J. van Moorsel, Terhi Hahl, Owen L. Petchey, Anne Ebeling, Nico Eisenhauer, Bernhard Schmid, and Cameron Wagg

**Table of Contents**

Page 2: **Appendix S1: FIG. S1.** Biomass stability and climate stability from 2003-2010.
Page 4: **Appendix S1: FIG. S2.** Aboveground community biomass over time.
Page 5: **Appendix S1: FIG. S3.** Effect sizes for fixed factors.
Page 6: **Appendix S1: FIG. S4.** Resistance, recovery, and resilience corrected for pre-flood biomass.
Page 7: **Appendix S1: FIG. S5.** Biodiversity–turnover relationship.
Page 8: **Appendix S1: TABLE S1.** ANOVA results for resistance, recovery, and resilience corrected for pre-flood biomass.
Page 9: **Appendix S1: TABLE S2.** ANOVA results for pre-flood stability and post-flood stability.
Page 10: **Appendix S1: TABLE S3.** ANOVA results for Bray-Curtis compositional turnover
Page 11: **Appendix S1: TABLE S4.** Differences between soil treatments in 2015.
Page 12: **Appendix S1: TABLE S5.** Species list.
Page 14: **Appendix S1: TABLE S6.** Overview seeds collected in Jena plots.
Page 16: **Appendix S1: Literature cited**

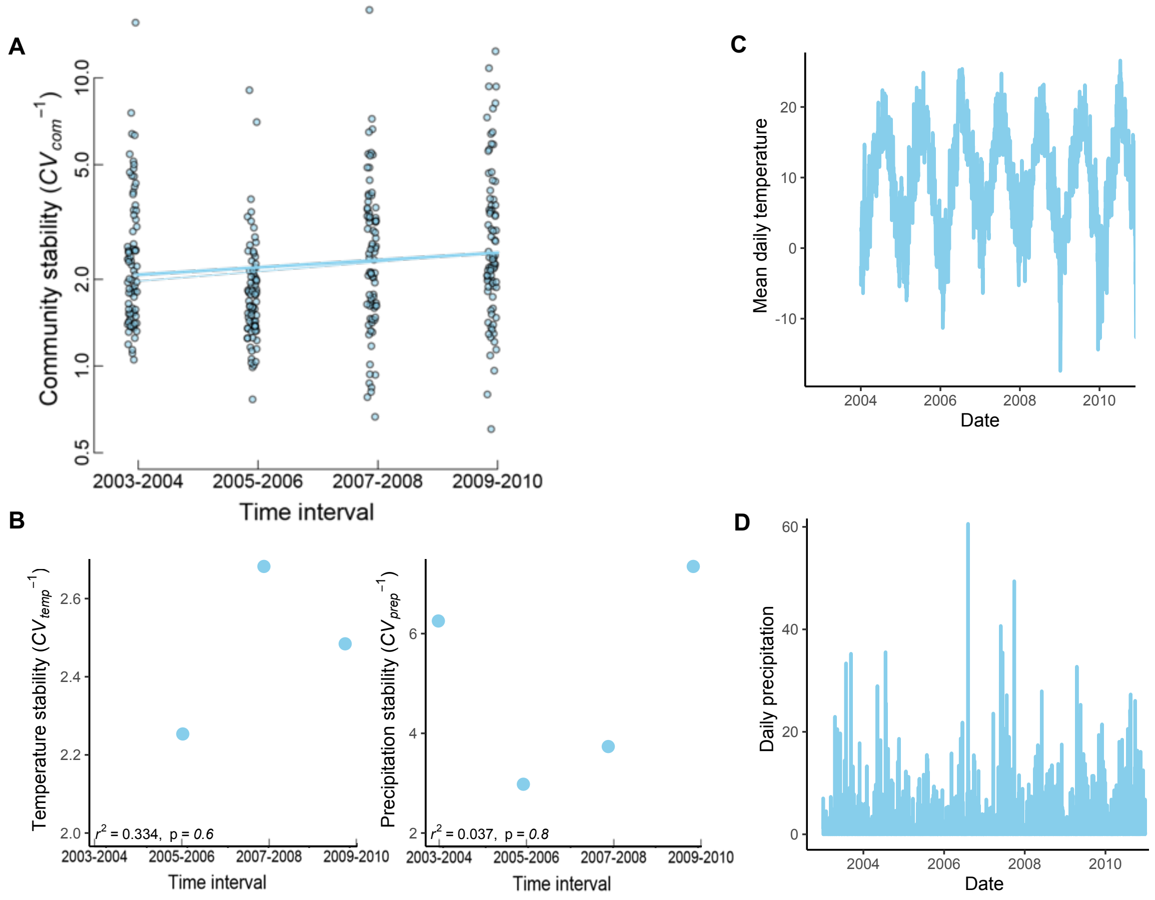


**Appendix S1: FIG. S1. Stability of community biomass and climate from 2003 to 2010.** (**A**) Combined intra- and inter-annual stability of experimental communities over the first 8 years in a grassland biodiversity experiment (Jena Experiment; species richness levels: 1, 2, 4, 8, 16, 60). The 8-year period was partitioned into four 2-year periods and within each stability was calculated for spring and summer harvests in year n and spring harvest in year n+1, corresponding to the same sequence of three harvests used in subsequent tests communities collected from the Jena Experiment in 2010 (selected communities) or re-established from seeds of the original supplier (naïve communities). Thick regression line includes three outliers outside the top margin of the plot ($P = 0.037$), thin line excludes these outliers ($P = 0.0018$). Changes in community biomass stability over time were also significantly correlated with precipitation stability ($P < 0.001$ when "precipitation stability" is fitted in the model instead of the term "time"). **(B)** Stability (inverse of the CV) over time for mean temperatures and precipitation in spring (March-May) and summer (June-August), times that correspond to the growth of biomass. The CV was calculated across three time points (spring year n, summer year n and spring year n+1). Temperature from the year 2003 is missing, which is why the first value appears in 2004. Note that the CV is the inverse of stability, thus lower values mean higher stability. Test statistics are shown in the figure. (**C**) Mean temperatures from 2003 to 2010. (**D**) Total daily precipitation from 2003 to 2010.

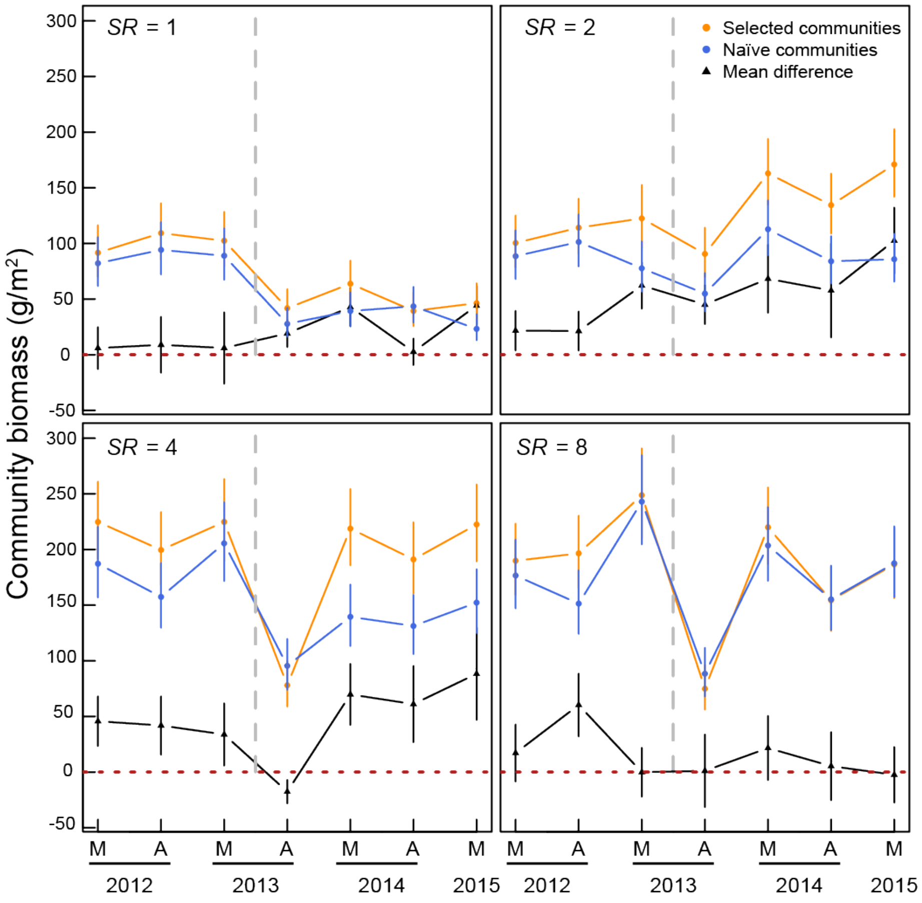


**Appendix S1: FIG. S2. Aboveground community biomass over time at four species richness levels (SR).** Selected and naïve plant communities and their mean difference are plotted with means and standard errors calculated from raw data. The dashed line indicates the flood event. M = May, A= August. For the calculation of resistance, resilience, and recovery, we averaged the community biomass in May 2012, August 2012, and May 2013 to obtain pre-flood biomass. We used the August 2013 biomass as our measure of biomass during the flood (even though we harvested several weeks after the water had receded). For post-flood biomass we averaged community biomasses from May 2014, August 2014, and May 2015.

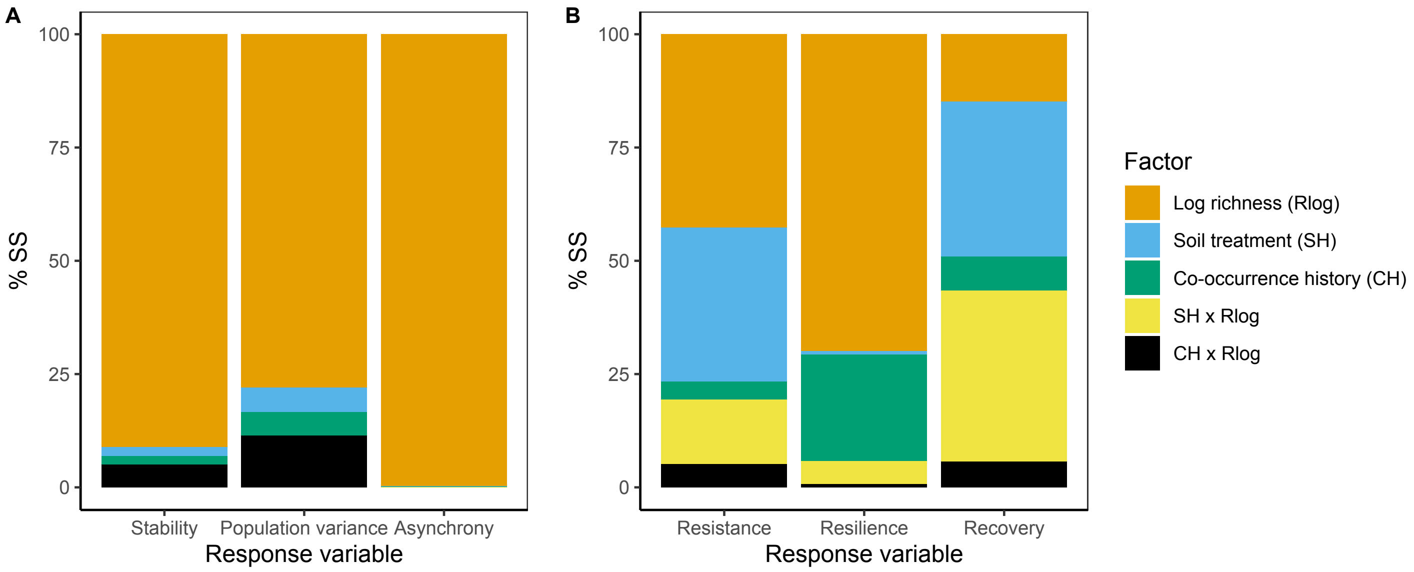


**Appendix S1: FIG. S3. Effect sizes (% SS) for fixed factors from a linear model. (A)** Asynchrony, population variance and community stability. **(B)** Recovery, resilience, and resistance**.** We used linear models to get % SS as effect sizes to compare relative explanatory power of the different fixed effects tested in the mixed models as done in hierarchical partitioning(Grömping 2006). Note that, due the almost fully orthogonal experimental design, % SS for different fitting sequences and results from linear and mixed models were nearly identical.

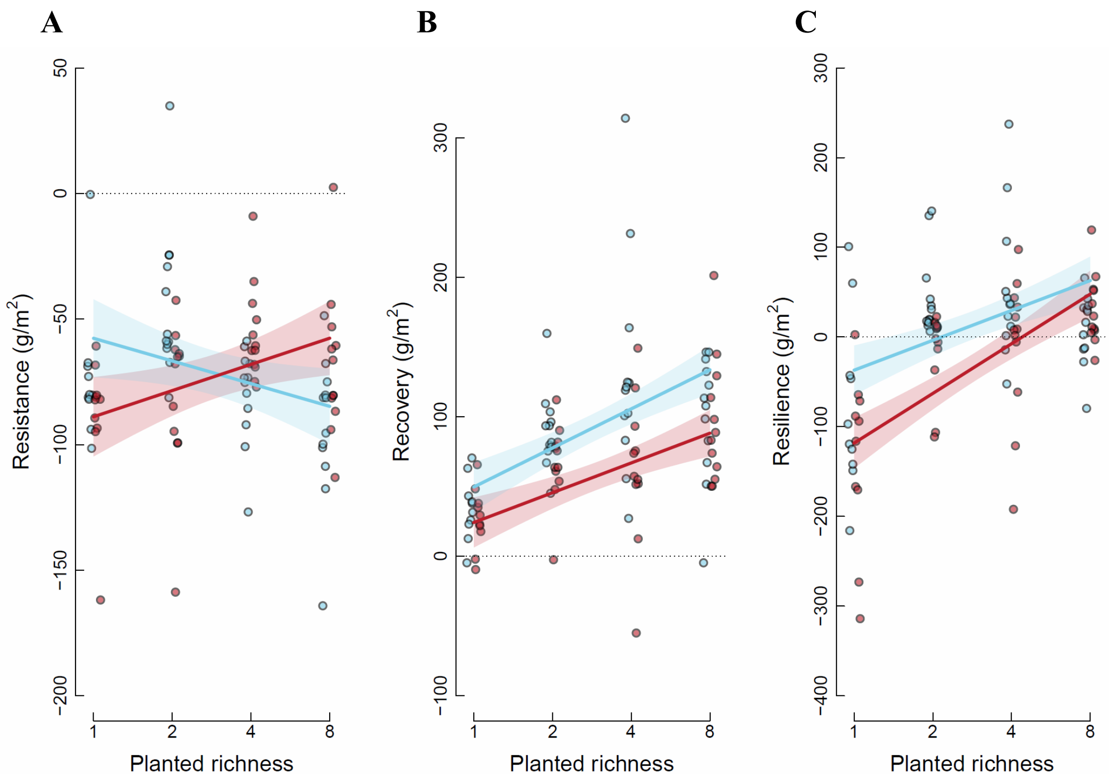


**Appendix S1: FIG. S4. Resistance, recovery, and resilience corrected for pre-flood biomass.** **(A)** Biodiversity–resistance relationships, **(B)** biodiversity–recovery relationships and **(C)** biodiversity–resilience relationships for selected (blue) and naïve communities (red). Colored bands indicate standard errors of predictions from mixed models as presented in Table S1. In contrast to Fig. 4 in the main text here the raw data were not only corrected for variation within diversity levels between plots and quadrats but also for variation in pre-flood biomass. Means across the three soil treatments are shown. The dashed line is drawn at 0 in each graph.

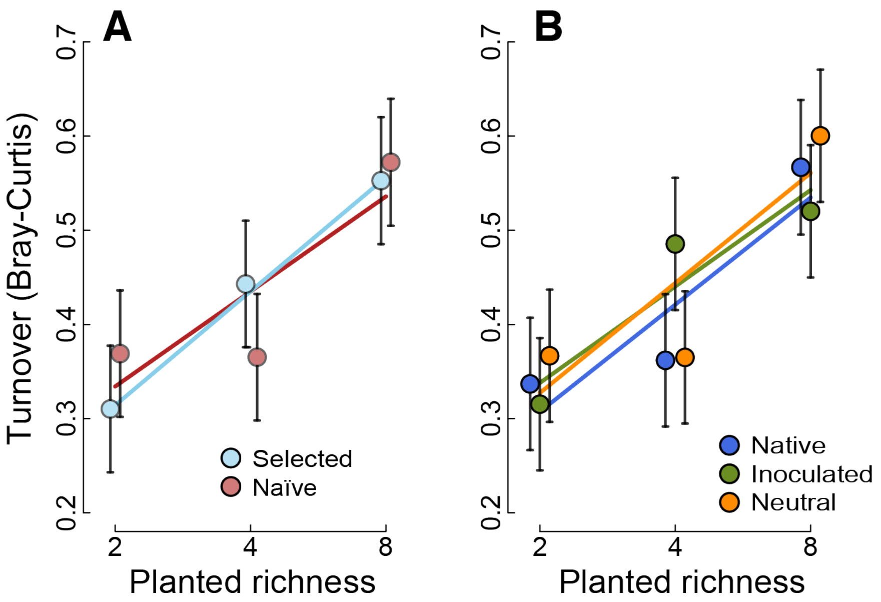


**Appendix S1: FIG. S5. The biodiversity–turnover relationship. (A)** Selected (blue) and naïve communities (red). **(B)** Home soil (blue), sterilized soil with native inoculum ("inoculated", green) and sterilized soil with neutral inoculum ("neutral", orange). Species compositional turnover was calculated between three pre- and three post-flood harvests. The species richness effect was significant but none of the other effects and none of the interactions were significant (see Appendix S1: Table S3). Shown are predicted means and standard errors.

**Appendix S1: TABLE S1.** Mixed-model ANOVA results for pre-flood biomass-corrected resistance, recovery, and resilience of community biomass. The effects of species richness (log scale), soil treatments and co-occurrence history on responses of community biomass to flooding were analyzed. In contrast to Table 2 (and the corresponding Fig. 4) in the main text, here the average of the three harvests before the flood (pre-flood productivity) was included as a covariate to account for the dependence of resistance, recovery, and resilience measures on the initial productivity. Bold italic text highlights significant effects.

| | | **Resistance** | | | **Recovery** | | | **Resilience** | | |
|---|---|---|---|---|---|---|---|---|---|---|
| Fixed terms | $DF_{num}$ | $DF_{den}$ | *F* | *P* | $DF_{den}$ | *F* | *P* | $DF_{den}$ | *F* | *P* |
| Pre-flood productivity | 1 | ***239.0*** | ***271.60*** | ***<0.001*** | ***199.7*** | ***17.05*** | ***<0.001*** | ***223.8*** | ***47.75*** | ***<0.001*** |
| Log richness ($R_{log}$) | 1 | 48.4 | 0.02 | 0.886 | ***49.6*** | ***9.53*** | ***0.003*** | ***49.2*** | ***11.50*** | ***0.001*** |
| Soil treatment (SH) | 2 | 96.2 | 0.41 | 0.668 | 95.2 | 0.13 | 0.877 | 96.4 | 1.50 | 0.229 |
| Co-occurrence history (CH) | 1 | 140.3 | 0.03 | 0.860 | ***140.0*** | ***11.65*** | ***<0.001*** | ***140.7*** | ***10.95*** | ***0.001*** |
| SH x $R_{log}$ | 2 | 90.3 | 1.23 | 0.296 | 89.5 | 2.39 | 0.097 | ***90.5*** | ***3.75*** | ***0.027*** |
| CH x $R_{log}$ | 1 | ***133.9*** | ***8.77*** | ***0.004*** | 134.4 | 0.48 | 0.491 | 134.6 | 2.84 | 0.094 |
| Random terms | *N* | *Var.* | *SE* | | *Var.* | *SE* | | *Var.* | *SE* | |
| Plot | 46 | 2124 | 643 | | 1887 | 717 | | 5971 | 1920 | |
| Plot x SH | 137 | 669 | 429 | | 112 | 756 | | 1265 | 1521 | |
| Residual | 274 | 3534 | 432 | | 7583 | 925 | | 14035 | 1711 | |

*Note*: $DF_{num}$ = numerator degrees of freedom, $DF_{den}$ = denominator degrees of freedom, *F* = variance ratio, *P* = probability of type-I error.

**Appendix S1: TABLE S2.** Mixed-model ANOVA results for log-transformed community stability for the three harvests before the flood event in late spring of June 2013 (pre-flood stability) and the three harvests after recovery from the flood event (post-flood stability). The effects of species richness (log scale), soil treatments, and co-occurrence history on the pre- and post-flood stability of community biomass were analyzed. Bold italic text highlights significant effects.

| | | Pre-flood stability | | | Post-flood stability | | |
|---|---|---|---|---|---|---|---|
| Fixed terms | $DF_{num}$ | $DF_{den}$ | *F* | *P* | $DF_{den}$ | *F* | *P* |
| Log richness ($R_{log}$) | 1 | 44.1 | 1.67 | 0.203 | ***43.9*** | ***13.89*** | ***<0.001*** |
| Soil treatment (SH) | 2 | 86.2 | 1.04 | 0.356 | 86.3 | 0.99 | 0.377 |
| Co-occurrence history (CH) | 1 | 133.1 | 1.50 | 0.222 | ***133.6*** | ***5.03*** | ***0.027*** |
| SH x $R_{log}$ | 2 | 87.9 | 2.26 | 0.110 | 87.1 | 0.28 | 0.754 |
| CH x $R_{log}$ | 1 | 134.5 | 2.86 | 0.093 | 134.1 | 0.10 | 0.749 |
| Random terms | *N* | *Var.* | *SE* | | *Var.* | *SE* | |
| Plot | 36 | 0.273 | 0.069 | | 0.092 | 0.032 | |
| Plot x SH | 107 | 0.008 | 0.027 | | -0.013 | 0.030 | |
| Residual | 214 | 0.267 | 0.033 | | 0.321 | 0.039 | |

*Note*: $DF_{num}$ = numerator degrees of freedom, $DF_{den}$ = denominator degrees of freedom, *F* = variance ratio, *P* = probability of type-I error.

**Appendix S1: TABLE S3.** Mixed-model ANOVA results for Bray-Curtis compositional turnover between three pre- and three post-flood harvests. The effects of species richness (log scale), soil treatments, and co-occurrence history on the compositional turnover were analyzed. Bold italic text highlights significant effects.

| | | **Turnover** | | |
|---|---|---|---|---|
| Fixed terms | $DF_{num}$ | $DF_{den}$ | *F* | *P* |
| Log richness ($R_{log}$) | 1 | ***34.0*** | ***6.25*** | ***0.017*** |
| Soil treatment (SH) | 2 | 67.1 | 0.30 | 0.744 |
| Co-occurrence history (CH) | 1 | 105.0 | 0.00 | 1.000 |
| SH x $R_{log}$ | 2 | 67.1 | 0.08 | 0.927 |
| CH x $R_{log}$ | 1 | 105.0 | 0.40 | 0.527 |
| Random terms | *N* | *Var.* $10^{-3}$ | *SE* $10^{-3}$ | |
| Plot | 36 | 41.29 | 11.55 | |
| Plot x SH | 107 | 0.00 | 4.00 | |
| Residual | 214 | 35.95 | 4.96 | |

**Appendix S1: TABLE S4. Analysis of soil-history treatments at the end of the experiment in October 2015.** Means and standard errors (SEMs) are given together with the *P*-values testing the significance of treatment effects in analyses of variance. SEMs were calculated with the raw data.

| | **Native soil** | | **Sterilized soil with native inoculum** | | **Sterilized soil with neutral inoculum** | | |
|---|---|---|---|---|---|---|---|
| Soil characteristics | Mean | SEM | Mean | SEM | Mean | SEM | Significance |
| Nitrate (ppm) | 7 | 0.26 | 5.7 | 0.26 | 5.5 | 0.25 | < 0.001 |
| Phosphorous (ppm) | 23.5 | 1.5 | 31.1 | 1.8 | 31 | 1.9 | < 0.001 |
| Microbial carbon | 626.5 | 16.1 | 451.8 | 14.2 | 442.3 | 14.6 | < 0.001 |
| Microbial nitrogen | 150.7 | 3.5 | 112.2 | 3.1 | 106.1 | 3.3 | < 0.001 |
| Bacterial richness (# 16S-OTUs) | 5230.4 | 71.1 | 4919.9 | 82 | 4822.5 | 92.1 | < 0.001 |
| Bacterial evenness | 0.889 | 8E-04 | 0.875 | 0.0007 | 0.864 | 0.00082 | < 0.001 |
| Fungal richness (# ITS-OTUs) | 774.8 | 17.9 | 765.7 | 17.6 | 765.9 | 19 | 0.1 |
| Fungal evenness | 0.879 | 0.002 | 0.885 | 0.0013 | 0.888 | 0.00148 | < 0.001 |

**Appendix S1: TABLE S5. Species list.** In the 47 experimental communities, a total of 49 species were grown in different community diversities and compositions. The eleven species occurring in monoculture are highlighted in bold. For species authorities and definition of functional groups see (Roscher et al. 2004). Biomass values are taken from small 3.5 x 3.5 m monoculture plots and represent yearly aboveground averages from 2003–2006 (Marquard et al. 2013).

| Species | Functional group | Life cycle | Self-incompatible (yes/no) | Biomass (g/m$^2$) |
|---|---|---|---|---|
| *Achillea millefolium* | herb | perennial | yes | 338.0 |
| *Ajuga reptans* | herb | perennial | no | 10.1 |
| *Alopecurus pratensis* | grass | perennial | no | 433.9 |
| *Anthoxanthum odoratum* | grass | perennial | no | 259.6 |
| *Arrhenatherum elatius* | grass | perennial | yes | 616.4 |
| *Avenula pubescens* | grass | perennial | yes | 422.6 |
| *Bromus erectus* | grass | perennial | yes | 675.5 |
| *Bromus hordeaceus* | grass | annual–biennial | no (mostly selfing) | 251.6 |
| ***Crepis biennis*** | herb | perennial | no | 326.4 |
| *Cynosurus cristatus* | grass | perennial | yes | 78.2 |
| *Dactylis glomerata* | grass | perennial | yes | 462.5 |
| *Daucus carota* | herb | biennial | yes | 376.9 |
| *Festuca pratensis* | grass | perennial | yes | 329.9 |
| ***Festuca rubra*** | grass | perennial | no | 334.7 |
| ***Galium mollugo*** | herb | annual | no | 438.1 |
| ***Geranium pratense*** | herb | perennial | no | 262.1 |
| *Glechoma hederacea* | herb | perennial | no | 92.8 |
| *Heracleum sphondylium* | herb | biennial–perennial | no | 180.0 |
| *Holcus lanatus* | grass | perennial | mostly yes | 500.7 |
| *Knautia arvensis* | herb | perennial | no | 644.4 |
| ***Lathyrus pratensis*** | legume | perennial | no | 357.8 |
| *Leontodon autumnalis* | herb | perennial | yes | 290.8 |
| *Leontodon hispidus* | herb | perennial | no | 331.8 |
| *Leucanthemum vulgare* | herb | perennial | yes | 445.6 |
| *Lotus corniculatus* | legume | perennial | mostly yes | 388.0 |
| *Luzula campestris* | grass | perennial | mostly yes | 0.1 |
| *Medicago lupulina* | legume | annual–perennial | no | 52.4 |
| *Medicago x varia* | legume | perennial | no | 815.9 |
| ***Onobrychis viciifolia*** | legume | perennial | no | 1290.5 |
| *Phleum pratense* | grass | perennial | mostly yes | 417.8 |
| ***Plantago lanceolata*** | herb | perennial | yes | 224.6 |
| *Plantago media* | herb | perennial | no | 420.8 |
| ***Poa pratensis*** | grass | perennial | no | 235.0 |
| *Poa trivialis* | grass | perennial | no | 164.7 |
| *Primula veris* | herb | perennial | yes | 168.1 |

| | | | | |
|---|---|---|---|---|
| ***Prunella vulgaris*** | herb | perennial | no | 222.3 |
| *Ranunculus acris* | herb | perennial | yes | 242.7 |
| *Ranunculus repens* | herb | perennial | yes | 132.4 |
| *Sanguisorba officinalis* | herb | perennial | no | 414.7 |
| *Taraxacum officinale* | herb | perennial | yes | 286.2 |
| *Trifolium campestre* | legume | annual | no | 8.9 |
| *Trifolium dubium* | legume | annual | yes? | 2.8 |
| *Trisetum flavescens* | grass | perennial | yes? | 422.6 |
| *Trifolium fragiferum* | legume | perennial | mostly yes | 143.1 |
| *Trifolium hybridum* | legume | perennial | mostly yes | 227.1 |
| *Trifolium pratense* | legume | perennial | yes | 353.1 |
| ***Trifolium repens*** | legume | perennial | yes | 361.4 |
| ***Veronica chamaedrys*** | herb | perennial | yes | 220.2 |
| *Vicia cracca* | legume | perennial | no | 93.2 |

**Appendix S1: TABLE S6. Overview of seeds collected in the Jena plots.** For those species that did not produce enough seeds in the experimental garden in Zurich, some additional seeds were collected directly in the Jena experimental plots. Shown are percentages of total seed weight with an origin of the Jena plots for each species in each experimental community.

| Plot | SR | Species | %seeds collected in Jena | Plot | SR | Species | %seeds collected in Jena | Plot | SR | Species | %seeds collected in Jena |
|---|---|---|---|---|---|---|---|---|---|---|---|
| **B1A01** | **16** | Pla lan | 21.4 | **B2A01** | **4** | Ant odo | 0.0 | **B3A04** | **8** | Alo pra | 0.0 |
| | | Lat pra | 0.0 | | | Pru vul | 0.0 | | | Cyn cri | 0.0 |
| | | Poa pra | 0.0 | | | Kna arv | 0.0 | | | Fes rub | 0.0 |
| | | Ger pra | 1.1 | | | Tri pra | 0.0 | | | Poa tri | 0.0 |
| **B1A02** | **8** | Alo pra | 29.7 | **B2A02** | **2** | Fes rub | 0.0 | | | Arr ela | 0.0 |
| | | Bro ere | 0.0 | | | Tri fla | 0.0 | | | Dac glo | 0.0 |
| | | Car pra | 0.0 | **B2A03** | **60** | Fes pra | 100.0 | | | Hol lan | 0.0 |
| | | Her sph | 0.0 | | | Fes rub | 0.0 | | | Tri fla | 0.0 |
| | | Fes rub | 0.0 | | | Pru vul | 0.0 | **B3A05** | **8** | Ant odo | 0.0 |
| | | Phl pra | 0.0 | | | Ver cha | 100.0 | | | Bro ere | 0.0 |
| | | Ran acr | 63.0 | | | Poa pra | 0.0 | | | Poa tri | 20.5 |
| | | San off | 0.0 | | | Pla lan | 100.0 | | | Ant syl | 100.0 |
| **B1A03** | **8** | Cyn cri | 0.0 | **B2A04** | **1** | Ger pra | 0.0 | | | Leu vul | 0.0 |
| | | Phl pra | 0.0 | **B2A05** | **1** | Fes pra | 0.0 | | | Lot cor | 5.0 |
| | | Gle hed | 0.0 | **B2A06** | **4** | Pla lan | 10.3 | | | Ono vic | 99.9 |
| | | Pri ver | 0.0 | | | Tar off | 0.0 | | | Tri hyb | 0.0 |
| | | Tri fla | 0.0 | | | Lat pra | 73.7 | **B3A06** | **1** | Fes rub | 53.0 |
| | | Ver cha | 0.0 | | | Med lup | 0.0 | **B3A07** | **8** | Bro hor | 0.0 |
| | | Lot cor | 0.0 | **B2A08** | **2** | Ran acr | 20.4 | | | Hol lan | 0.0 |
| few seed | | Med lup | 0.0 | | | Tri cam | 0.0 | | | Pri ver | 0.0 |
| **B1A04** | **4** | Fes pra | 0.0 | **B2A09** | **4** | Aju rep | 0.0 | | | Ran rep | 100.0 |
| | | Pla lan | 33.0 | | | Pla lan | 4.6 | | | Her sph | 0.0 |
| | | Cam pat | 0.0 | | | Pri ver | 0.0 | | | Leu vul | 0.0 |
| | | Ono vic | 0.0 | | | Pru vul | 3.6 | | | Med lup | 0.0 |
| **B1A05** | **2** | Med lup | 0.0 | **B2A12** | **8** | Ant syl | 0.0 | | | Ono vic | 82.7 |
| | | Ono vic | 0.0 | | | Ger pra | 0.0 | **B3A08** | **2** | Dac glo | 0.0 |
| **B1A07** | **2** | Ran acr | 17.2 | | | Kna arv | 52.6 | | | Fes pra | 0.0 |
| | | San off | 0.0 | | | Ran acr | 4.3 | **B3A09** | **16** | Fes pra | 94.0 |
| **B1A11** | **16** | Ger pra | 0.0 | | | Gal mol | 0.0 | | | Fes rub | 0.0 |
| | | Cre bie | 10.1 | | | Her sph | 0.0 | | | Poa pra | 0.0 |
| | | Gal mol | 0.0 | | | Leu vul | 0.0 | **B3A11** | **4** | Bro ere | 0.0 |
| **B1A12** | **8** | Lat pra | 0.0 | | | San off | 0.0 | | | Poa tri | 0.0 |
| | | Med var | 0.0 | **B2A13** | **1** | Pla lan | 1.2 | | | Pla lan | 6.7 |
| few seed | | Tri cam | 0.0 | **B2A14** | **8** | Luz cam | 0.0 | | | Pru vul | 2.1 |
| | | Tri hyb | 0.0 | | | Phl pra | 0.0 | **B3A12** | **1** | Lat pra | 30.1 |
| | | Med lup | 0.0 | | | Leo his | 0.0 | **B3A13** | **4** | Alo pra | 0.0 |
| | | Ono vic | 81.2 | | | Ver cha | 0.0 | | | Bro ere | 96.3 |
| | | Tri dub | 0.0 | | | Kna arv | 79.9 | | | Ant odo | 0.0 |
| | | Tri pra | 0.0 | | | San off | 0.0 | | | Poa tri | 0.0 |
| **B1A13** | **4** | Lot cor | 0.0 | | | Tri dub | 0.0 | **B3A17** | **1** | Ver cha | 30.0 |
| | | Med var | 0.0 | | | Tri hyb | 5.5 | **B3A19** | **2** | Tri fla | 0.0 |
| | | Ono vic | 0.0 | **B2A15** | **1** | Ono vic | 48.1 | | | Tar off | 0.0 |
| | | Med lup | 0.0 | **B2A16** | **4** | Leo aut | 0.0 | **B3A21** | **2** | Lot cor | 0.5 |

| | | | |
|---|---|---|---|
| **B1A14** | **8** | Luz cam | 0.0 |
| | | Tri fla | 0.0 |
| | | Leo his | 0.0 |
| | | Pla lan | 28.4 |
| | | Ant syl | 0.0 |
| | | Dau car | 0.0 |
| | | Tri cam | 8.4 |
| | | Tri fra | 0.0 |
| **B1A15** | **1** | Cre bie | 0.0 |
| **B1A16** | **2** | Poa pra | 0.0 |
| | | Pla lan | 6.0 |
| **B1A17** | **2** | Alo pra | 46.6 |
| | | Dau car | 0.0 |
| **B1A18** | **1** | Pru vul | 2.4 |
| **B1A19** | **4** | Arr ela | 31.8 |
| | | Luz cam | 0.0 |
| | | Pru vul | 0.0 |
| | | Cam pat | 0.0 |
| **B1A21** | **4** | Fes pra | 0.0 |
| | | Luz cam | 0.0 |
| | | Ach mil | 0.0 |
| | | Cre bie | 0.0 |
| **B1A22** | **60** | Fes pra | 0.0 |
| | | Fes rub | 0.0 |
| | | Pru vul | 0.0 |
| | | Ver cha | 0.0 |
| | | Ger pra | 0.0 |
| | | Poa pra | 0.0 |
| | | Pla lan | 8.1 |
| | | Pla med | 9.6 |
| | | Kna arv | 80.4 |
| | | Vic cra | 0.0 |
| **B2A17** | **8** | Gle hed | 0.0 |
| | | Pla med | 25.2 |
| | | Leo aut | 0.0 |
| | | Tar off | 0.0 |
| | | Lat pra | 0.0 |
| | | Vic cra | 47.9 |
| | | Tri cam | 0.0 |
| | | Tri fra | 0.0 |
| **B2A18** | **16** | Poa pra | 0.0 |
| | | Ger pra | 0.0 |
| | | Tri rep | 18.5 |
| **B2A19** | **2** | Pla med | 23.9 |
| | | Tar off | 0.0 |
| **B2A20** | **2** | Pla lan | 8.0 |
| | | Tri dub | 0.0 |
| **B2A21** | **8** | Leo his | 23.0 |
| | | Pla med | 49.7 |
| | | Cre bie | 0.0 |
| | | Gal mol | 0.0 |
| | | Lot cor | 69.9 |
| | | Med lup | 0.0 |
| | | San off | 65.1 |
| | | Ono vic | 92.1 |
| **B3A01** | **1** | Gal mol | 92.5 |
| **B3A02** | **2** | Fes pra | 0.0 |
| | | Car car | 0.0 |
| **B3A03** | **4** | Phl pra | 0.0 |
| | | Pla med | 55.3 |
| | | Tri hyb | 0.0 |
| | | Vic cra | 25.1 |
| | | Tri pra | 0.0 |
| **B3A22** | **16** | Fes rub | 0.0 |
| | | Ver cha | 0.0 |
| | | Cre bie | 0.7 |
| | | Ger pra | 41.6 |
| | | Gal mol | 99.7 |
| | | Pla lan | 100.0 |
| | | Ono vic | 0.0 |
| **B4A06** | **8** | Pru vul | 0.0 |
| | | Ver cha | 0.0 |
| **B4A08** | **8** | Ant odo | 0.0 |
| | | Bro hor | 0.0 |
| | | Ave pub | 0.0 |
| | | Fes rub | 0.0 |
| | | Aju rep | 0.0 |
| | | Tar off | 0.0 |
| | | Pla lan | 22.1 |
| | | Ver cha | 100.0 |
| **B4A09** | **1** | Tri rep | 0.0 |
| **B4A12** | **1** | Poa pra | 45.1 |
| **B4A18** | **16** | Ver cha | 78.1 |
| | | Cre bie | 0.0 |
| | | Lat pra | 83.8 |
| | | Ono vic | 97.5 |
| **B4A22** | **4** | Cam pat | 0.0 |
| | | Ger pra | 0.0 |
| | | Car pra | 0.0 |
| | | Kna arv | 0.0 |

### Appendix S1: LITERATURE CITED

Grömping, U. 2006. Relative Importance for Linear Regression in *R* : The Package **relaimpo**. Journal of Statistical Software 17.

Marquard, E., B. Schmid, C. Roscher, E. De Luca, K. Nadrowski, W. W. Weisser, and A. Weigelt. 2013. Changes in the Abundance of Grassland Species in Monocultures versus Mixtures and Their Relation to Biodiversity Effects. PLoS ONE 8:e75599.

Roscher, C., J. Schumacher, J. Baade, W. Wilcke, G. Gleixner, W. W. Weisser, B. Schmid, and E.-D. Schulze. 2004. The role of biodiversity for element cycling and trophic interactions: an experimental approach in a grassland community. Basic and Applied Ecology 5:107–121.
